# H2AX C-Terminal Dipeptide Truncation: A Master Switch of the DNA Damage Response

**DOI:** 10.64898/2026.05.07.723272

**Authors:** Faith M. Joseph, Matthew V. Holt, John M. Jerome, Linda Zhang, Ashley G. Boice, Patricia D. Castro, Sofía I. Aramburu, Ruhee Dere, Susan M. Rosenberg, David R. Rowley, Nicolas L. Young

## Abstract

Phosphorylation of histone H2AX at serine 139 (γH2AX) by ATM/ATR kinases is a central marker of the DNA damage response (DDR), widely used to detect DNA double-strand breaks. However, the molecular basis for tissue- and context-specific variation in γH2AX signaling remains poorly defined. Here we discover a post-translational truncation of H2AX, catalyzed by lysine demethylase 4A (KDM4A), which removes two C-terminal amino acids critical for ATM/ATR-dependent phosphorylation. This truncation renders H2AX refractory to γH2AX formation, effectively bypassing canonical DDR signaling. Truncated H2AX accumulates in select cell lines, primary cells, solid tumors, and normal tissues. Genetic knockdown or pharmacologic inhibition of KDM4A reduces H2AX truncation, restores γH2AX induction, and enhances DNA repair capacity. Conversely, KDM4A overexpression promotes H2AX truncation, impairs γH2AX signaling, and exacerbates DNA damage accumulation. This previously unrecognized regulatory axis implicates KDM4A catalyzed H2AX truncation as a superseding mechanism that represses the canonical DDR and disrupts the correlation between γH2AX and DNA damage. This dioxygenase-based protease mechanism represents a new class of proteases and is the first example of c-terminal dipeptide protein truncation. This discovery has broad implications in the basic science of genome maintenance, wound healing, cancer, combinatorial therapy, precision medicine, and technologies such as gene editing.

**Graphical Abstract:** 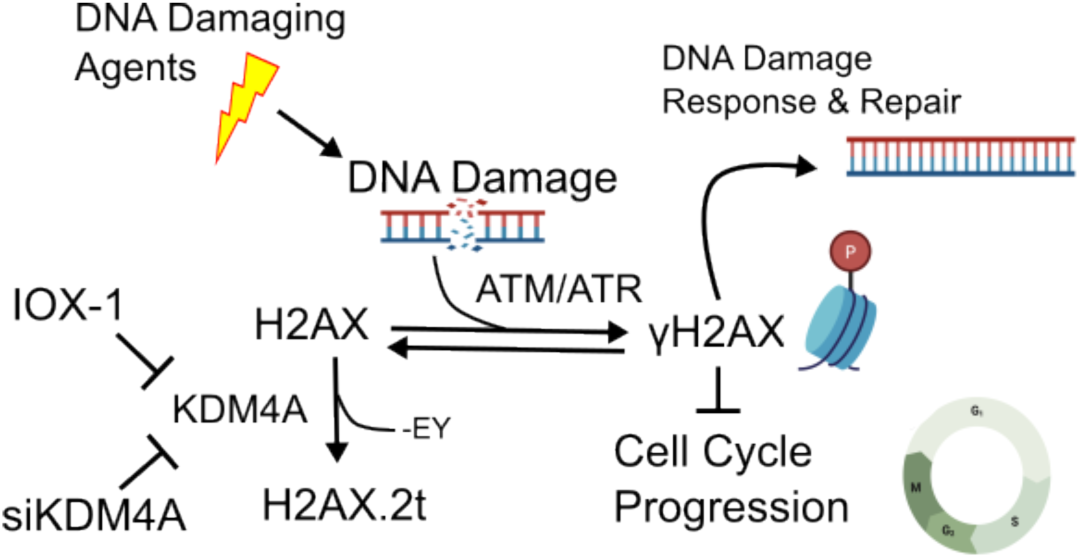

## Introduction

Histone proteins compact DNA into nucleosomes and govern access to the eukaryotic genome. Nucleosomes are composed of two copies of histone families H2A, H2B, H3, and H4, occasionally accompanied by the linker histone family H1. Through their post-translational modifications (PTMs), histones assume diverse roles in gene expression, chromosome segregation, DNA repair, and other fundamental chromosomal processes^1^. Each histone family is comprised of multiple genes with varying divergence in DNA and protein sequences. These many related sequences are prone to different modifications and carry out different functions^2^.

The prevailing approaches used to study histones primarily target known attributes and are prone to overlooking clipping events and their functions. The histone variant H2AX was discovered in 1980 and is well-studied due to its pivotal role in the DNA damage response. Phosphorylation of H2AX at serine 139, close to the C-terminus (-S_139_QEY), initiates DNA repair^3^. Current dogma highlights S139 phosphorylation of H2AX (γH2AX) by the PI3K kinases ATM/ATR as an early signal of the DNA damage response. The γH2AX signal initiates downstream mechanisms of repair, dependent on break type and severity^4–8^. Thus, γH2AX is used extensively as a marker of DNA damage for both basic science and clinical applications.

We discover here that the γH2AX signal is prone to a superseding regulatory mechanism that substantially weakens the positive correlation between γH2AX and DNA damage and allows for the accumulation of DNA damage that is not accompanied by sufficient γH2A for a robust DNA damage response. Substantial portions of H2AX molecules in a cell are often present in an enzymatically truncated and non-functional form that lacks the last two amino acids. Specifically, the truncated form consists of the first 140 amino acids, excluding the initiator methionine and including an N-terminal acetylation that is constitutively present on all H2AX molecules. This is denoted as {H2AX_[1-140]_Nα-ac} per N^3^ nomenclature^9^; however, we refer to this as H2AX.2t throughout for brevity and clarity. Because this difference in sequence is small, H2AX.2t has previously escaped detection. We detected this event only because we routinely study intact histones by top down proteomics to understand chromatin biology with proteoform-specificity^10–16^. The well-established importance of the C-terminus of H2AX particularly in genome stability prompted further inquiry. Here we also reveal the mechanism, function, and biological significance of H2AX truncation in the DDR and repair process.

Protein post-translational modifications (PTM) are an essential mechanism of regulation of protein function and transduction of signals. This includes reversible addition of chemical modifications, such as H2AXS139ph; however, post-translational removal of amino acids or peptides from the N- or C-termini can serve a similar function. Protein termini often mediate important protein-protein interactions and a small change in the final amino acid sequence can dramatically affect function. Some post-translational proteolysis events are even reversible, such as the proteolytic processing and re-ligation of tyrosine from the -GEEY motif at the unstructured C-terminus of alpha-tubulin^17^. Proteolysis of histones is an established but understudied means of chromatin regulation^18,19^.

We show here that lysine demethylase 4A (KDM4A) catalyzes the proteolysis of the two C-terminal amino acids on H2AX. The loss of the terminal EY dipeptide evidently precludes ADP-ribosylation of E141 and phosphorylation of Y142, signals for base excision repair and apoptosis respectively^20^. Despite S139 being retained, H2AX.2t is refractory to S139 phosphorylation. H2AX.2t is found in patient tumor sections, primary cells, and immortalized cell lines. The abundance of H2AX.2t varies between cellular contexts but is tightly consistent within each cell line. H2AX.2t is absent from some cells but represents as much as 50% of total H2AX. H2AX.2t is reduced by KDM chemical inhibition or KDM4A knockdown, which reestablishes downstream γH2AX signaling and restores effective DNA damage repair. Many cells that are low in KDM4A have undetectable levels of H2AX.2t. Conversely, overexpression of KDM4A in these cells results in a dramatic increase in H2AX.2t. KDM4A directly acts on the H2AX C-Terminal peptide and removes the last two amino acids in vitro. This KDM4A dependent reduction in γH2AX signaling decreases DNA damage repair, resulting in an accumulation of damage. We support these conclusions with multiple lines of evidence using orthogonal methods, performed in multiple labs. Taken together, this evidence reveals that H2AX truncation is potentially an important regulator of the DDR that supersedes the known signaling pathway.

## Results

### Histone H2AX is commonly truncated by two amino acids on its C-terminus

The human genome contains about 29 genes in the Histone H2A family and these express around 19 distinct H2A protein sequences and many more proteoforms. Due to the overlap in amino acid composition, sequences of the same histone family are often indistinguishable by common methods. We specialize in using top-down mass spectrometry (long read protein sequencing of intact proteins) to overcome sequence ambiguities and study variant specific histone PTMs and proteoforms. By top-down proteomic analysis of the histone H2A family, we discovered that histone H2AX is truncated by two amino acids on its C-terminus and H2AX.2t is often similarly as abundant as H2AX. H2AX.2t was first identified as an unassigned peak in our H2AX spectra 200 Da lower in mass with a similar but distinct retention time to the full-length H2AX peak (Fig. 1a). The prevalence of the unexpected population varies between cell lines but is consistently expressed within each of the cell lines tested regardless of sample processing and mass spectrometry parameters. Amino acid sequences of both the unidentified and expected peak of H2AX are identical in sequence until the last two C-terminal amino acids, “EY” (Fig. 1b-c). This reveals a novel form of H2AX existing in conjunction with the known H2AX sequence.

**Fig. 1.**
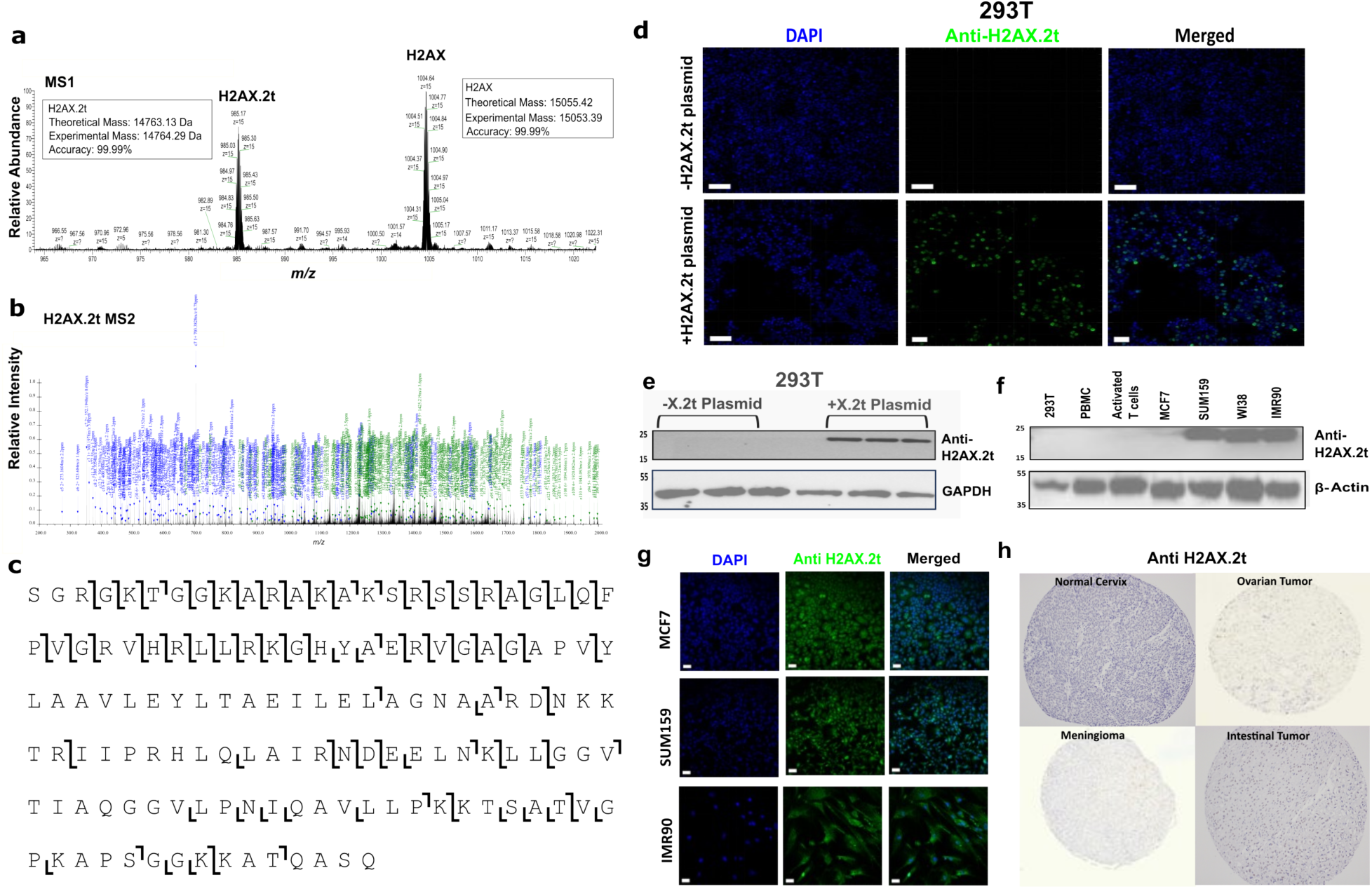
Histone H2AX is commonly truncated on its c-terminus by two amino acids in many cell lines, as well as normal and cancerous patient tissues. **a**, An MS1 spectrum of an H2A fraction extracted from SUM159 cells averaged from 48.68-49.02 minutes. We observe precursor masses consistent with full length histone H2AX and a shortened version of H2AX truncated by two amino acids (H2AX.2t). **b,** MS2 spectrum of the precursor 869.3224 *m/z* matching the expected fragments for H2AX.2t. An expanded version can be found in Supplementary Fig. 6. **c,** An ion map of H2AX.2t from the MS2 in Fig. 1b. This reveals sequence coverage that confirms the loss of the two C-terminal amino acids (EY) from H2AX. An ion map of full length H2AX can be found in Supplementary Fig. 1. **d,** Immunofluorescence (IF) of HEK293T cells with and without H2AX.2t plasmid demonstrates successful transfection, expression, and specific detection of H2AX.2t in cells. (H2AX.2t: #PP145, AF488: #22C0617) **e,** Western blot of H2AX.2t in HEK293T cells +/- plasmid demonstrates specific detection and confirms the results of Fig. 1d. (H2AX.2t plasmid: #60738, H2AX.2t: #PP9208, GAPDH: #14C10) **f,** Western blot analysis reveals varied expression levels of truncated H2AX in several cell lines, including primary fibroblasts. (H2AX.2t: #PWP201, β-Actin: #4967S) **g,** Immunofluorescence (IF) confirms the results in Fig. 1f and shows that H2AX.2t is primarily nuclear localized. (H2AX.2t: #PP145, AF488: #22C0617) **h,** Example TMA immunohistochemistry images reveal H2AX.2t is present in both cancerous and normal human tissues. Of 396 primarily tumor samples tested, about 5% are positive for H2AX.2t expression. H2AX.2t is more prominent in cancers from certain tissues such as breast. It is also intriguingly found in normal cervical tissue but absent in the cervical cancers tested. (H2AX.2t: #PWP9119) A summary of tumor and tissue results can be found in Supplementary Table 1.

### Histone H2AZ.1 but not H2AZ.2 is also truncated by two amino acids on its C-terminus

We also discovered that histone H2AZ.1, but not H2AZ.2, is commonly truncated by two amino acids on its C-terminus (Supplementary Fig. 2). H2AZ dysregulation is common in many cancers and like H2AX, the c-terminus of H2AZ is important to its gene regulatory function^21^. H2AZ has limited sequence homology with H2AX, and their c-terminal sequences are unrelated. H2AZ.1 and H2AZ.2 differ by three amino acids, with the c-terminal amino acid being one of these differences. This truncation at the penultimate amino acid is not observed on any other H2A sequences. While we do not focus on the mechanism and function of H2AZ truncation here, the two amino acid processing we observe on H2AX appears to be a conserved sequence-specific mechanism that also regulates other histone H2A variant-specific processes.

### H2AX.2t is detected in some normal tissues, tumors, primary, and immortalized cell lines

To complement our mass spectrometry discovery of H2AX.2t and enable further studies, we developed an H2AX.2t specific antibody (Fig. 1d-f). To validate the antibody, we used an H2AX.2t plasmid to express the truncated H2AX variant in HEK293T cells, a context where we only observe full length H2AX by mass spectrometry. As expected, we find that without plasmid transfection, there is no H2AX.2t signal in HEK293T cells by western and IF. Upon transfection of the truncated histone variant gene, we detect a signal for H2AX.2t (Fig. 1d-e). Thus, the antibody is specific to H2AX.2t over H2AX. We then used this antibody to confirm the presence of H2AX.2t by IF and western in the cell lines where it is observed by mass spectrometry.

With the H2AX.2t-specific antibody, we assessed multiple cell lines, cell types, and patient tissue sections. We detect H2AX.2t by western in SUM159 and MCF7 breast cancer cells; and normal fibroblast cell lines IMR90 and WI38. It is not detected in HEK293T cells, activated T cells, or peripheral blood mononuclear cells (PBMCs) (Fig. 1f). These results suggest that it is a normal regulatory mechanism active in some proliferative primary cells and appropriated by some cancers. Next, we used this antibody for immunofluorescence (IF) to show that H2AX.2t is localized to the nucleus similar to full length H2AX. In order of most to least intensity, SUM159, IMR90, and MCF7 cells express H2AX.2t and it is mostly nuclear localized (Fig. 1g). Partial cytoplasmic staining is observed in some cells, consistent with prior observation of context-dependent cytoplasmic localization of full length H2AX^22^. Lastly, we stained tissue microarray (TMA) slides of 396 patient samples which include various tumor types and stages as well as normal patient tissue. H2AX.2t is detected in around 1 in 20 tumors (Fig. 1h). It is also observed in some normal patient tissue sections, such as normal cervix, consistent with our observations in some cultured primary cells. These results provide further evidence that this novel variant is prevalent and is not specific to tumor cells alone.

### Jumonji C domain inhibitor N-Octyl IOX-1 attenuates H2AX.2t abundance

After eliminating other possibilities, we hypothesized that known epigenetic enzymes may mediate this truncation event. A small-scale screen identified only one treatment that reduced H2AX.2t abundance and did so dramatically. The N-octyl ester derivative of 5-carboxy-8-hydroxyquinoline (N-Octyl IOX-1) is a potent pan jumonji C domain inhibitor. Jumonji C domains are the dioxygenase catalytic domains of class two lysine demethylases (KDMs) and typically activate oxygen to remove methyl groups from methylated lysines. These enzymes have been shown in at least one previous instance to act as a histone protein hydrolase; however, in a methyllysine dependent manner^23^. N-Octyl IOX-1 reduces the abundance of H2AX.2t by mass spectrometry, western blot, and IF (Fig. 2a-c)^24^. SUM159 cells were treated with 4µM of N-Octyl IOX-1 for 24 hours and H2AX.2t abundance was quantified. Because N-Octyl IOX-1 reduces the abundance of H2AX.2t, we hypothesized that the dioxygenase function of class two KDMs is responsible for generating H2AX.2t. However, the effects of N-Octyl IOX-1 are not specific to any particular class 2 KDM.

**Fig. 2.**
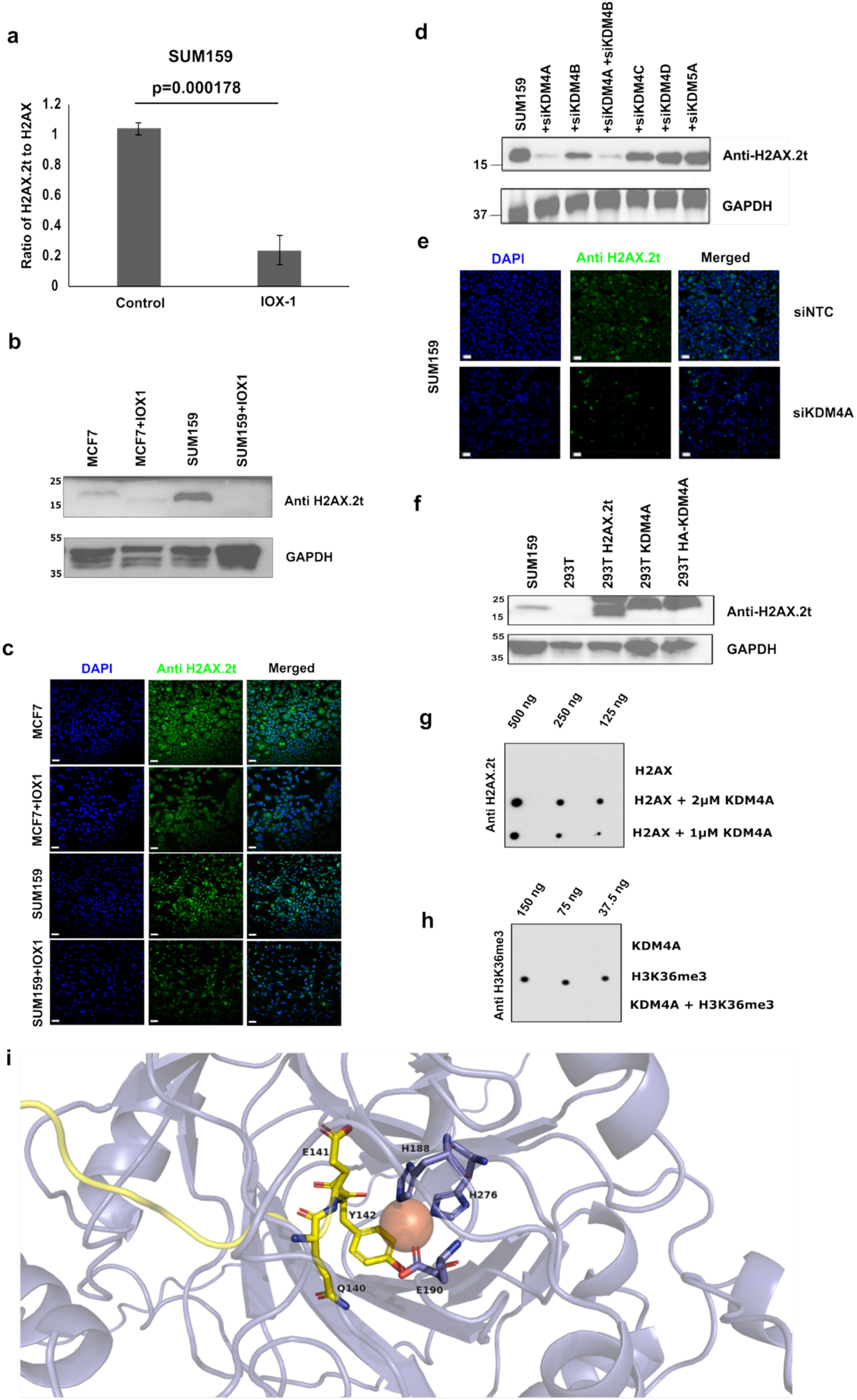
KDM4A catalyzes the sequence specific proteolysis of two c-terminal amino acids from H2AX. **a**, A 24-hour 4µM N-Octyl IOX-1 treatment significantly reduces the abundance of H2AX.2t as quantified by top-down mass spectrometry, (N=3). **b**, N-Octyl IOX-1 treatment reduces H2AX.2t in SUM159 and MCF7 cells by anti-H2AX.2t western blot. (H2AX.2t: #PWP201, GAPDH: #14C10) **c**, IF images of SUM159 and MCF7 cell lines show reduced nuclear expression of H2AX.2t with N-Octyl IOX-1 treatment (+IOX-1). All scale bars are 50µm. (H2AX.2t: #PWP201, AF488: #22C0617) **d**, Western blot of anti-H2AX.2t in SUM159 cells show a significant reduction in H2AX.2t upon KDM4A knockdown. (H2AX.2t: #PWP201, GAPDH: #14C10) **e**, H2AX.2t is detected in SUM159 cells but not detected in HEK293T cells. Direct expression of the H2AX.2t plasmid (Cat # 60738) in HEK293T cells exhibits strong H2AX.2t signal. Overexpression of two different KDM4A constructs (KDM4A = Full length KDM4A, HA-KDM4A = HA and FLAG tagged full length KDM4A) in HEK293T cells also increases the abundance of H2AX.2t. This indicates that KDM4A overexpression results in increased truncation of H2AX. (H2AX.2t: #PWP9208, GAPDH: #14C10) **f**, Knockdown of KDM4A in SUM159 cells reduces H2AX.2t by IF (H2AX.2t: #PP145, AF488: #22C0617). **g**, An in vitro activity assay performed for 2 hours with recombinant KDM4A and the C-terminal peptide of H2AX. Activity is seen with both 1µM and 2µM KDM4A and an approximately 2x difference in signal is observed. Serial dilutions of the reaction products (500, 250, and 125 ng) were loaded onto the dot blot as indicated. (H2AX.2t: PWP9119) **h**, A control in vitro activity assay for the canonical histone H3K36 demethylase activity of KDM4A under similar conditions. Serial dilutions of the reaction products (150 ng, 75 ng, and 37.5 ng) were loaded onto the dot blot as indicated. (H3K36me3: #31580) **i,** Alphafold predicts that the c-terminal tyrosine of H2AX (Y142) engages the catalytic iron of the jumonji domain of KDM4A through a cation-π interaction. In this configuration the proteolyzed peptide bond between H2AX Q140 and E141 is held near the site of catalytic dioxygenase activity.

### KDM4A expression modulates H2AX.2t abundance

The finding that N-Octyl IOX-1 reduces H2AX.2t led us to investigate the specific KDM implicated in H2AX proteolysis^25^. We hypothesized that KDM4A is the most likely candidate due to its strong connection to genome instability and cancer^26–28^.

To test our KDM4A hypothesis, we tested all six KDM4 members and other representative KDMs. For 48 hours, we transfected SUM159 and MCF7 cells with siRNAs targeting KDMs 4A, 4B, 4C, 4D, 5A and 6B. We also included a condition knocking down KDM4A and B in the same sample because they are close homologs that are often concurrently overexpressed in cancer. KDM4A depletion almost completely abolishes H2AX.2t in both cell lines (MCF7 data not shown) (Fig. 2d). The reduction of H2AX.2t by siKDM4A is similarly as dramatic when observed by IF and remains nuclear localized (Fig. 2e) and is quantified in Supplementary Fig. 4. KDM4B depletion only slightly reduces H2AX.2t. Co-transfection of siKDM4A and B, reduces H2AX.2t levels, but a more prominent reduction is seen with transfection of siKDM4A alone.

Next, we sought to test the converse implication of our model with an overexpression assay. Untreated HEK293T cells display no expression of H2AX.2t observable by western or IF (Fig. 1d-f). Overexpression of KDM4A in these cells results in H2AX.2t levels greater than naturally found in SUM159 cells and is comparable to directly overexpressing an H2AX.2t plasmid (Fig. 2f).

### KDM4A directly catalyzes the truncation of H2AX to H2AX.2t in vitro

The evidence above strongly implicates KDM4A in the proteolytic cleavage of the last two amino acids of H2AX. The suggested mechanism makes sense in the context of well-established biology: H2AX is a central hub of the DDR and KDM4A promotes genome instability. Collectively, the evidence thus far is sufficient to establish a strong functional relationship with profound implications. However, these in vivo experiments do not preclude indirect mechanisms, such as KDM4A demethylase activity activating another enzyme. There is only one prior example in the literature suggesting that KDM proteins and their JMJC domains can function as proteases. There are no known enzymatic mechanisms of cleaving a dipeptide from the c-terminus of proteins; however, such dipeptide losses are very commonly observed in terminomics experiments using chemoenzymatic labeling^29,30^. Thus, we next sought to establish the feasibility of a direct mechanism in an isolated in vitro system.

Recombinant KDM4A is sufficient to cleave the last two amino acids of the H2AX (CKATQASQEY) C-terminal peptide (Fig. 2g-h). The H2AX peptide alone does not react with the anti-H2AX.2t antibody. H2AX C-terminal peptide + 2µM KDM4A and cofactors, robustly cleave the peptide. A dilution series of these reaction conditions result in a proportional reduction in detected product. Reducing the KDM4A concentration from 2µM to 1µM KDM4A, while holding other reaction components constant, results in a 2-fold reduction in truncated peptide.

Structural modeling of full length H2AX and KDM4A using AlphaFold indicates that an interaction between the C-terminus of H2AX and the Jumonji domains of KDM4A is likely (Fig. 2i; Supplementary Fig. 3)^31^. The modeled structure suggests a mechanism by which Y142 plays a crucial role by engaging with the catalytic iron through a cation-π interaction. This results in the presentation of the Q140 and E141 peptide bond with uninhibited access to the iron catalyzed dioxygenase activity of KDM4A. The Y142 dependence of the suggested mechanism also explains the observed sequence specificity. Consistent with this prediction, the removal of the JMJC catalytic iron from the 1µM KDM4A conditions results in trace production of truncated peptide only at the highest concentration (Supplementary Fig. 3e).

Collectively, these in vitro experiments support a direct mechanism by which KDM4A uses its jumonji dioxygenase domain to effect proteolysis, possibly through an activated water intermediate. This implies that the KDM family is potentially a new class of dual function proteases. However, we primarily seek here only to establish: 1) that H2AX is truncated by two amino acids in some cellular contexts; 2) that the tight functional relationship observed between KDM4A and H2AX.2t is feasible. This is necessary to next address the function and biological significance of this discovery.

### H2AX.2t is refractory to phosphorylation by ATM/ATR at Serine 139

Because γH2AX is fundamental to genome stability we endeavored to understand the function of H2AX.2t in the DNA damage response and repair system. It is apparent that the loss of the terminal EY precludes ADP-ribosylation of E141, a signal for base excision repair, and Y142 phosphorylation, a signal for apoptosis. However, truncation of H2AX preserves the crucial S139 residue but this does not mean it is prone to phosphorylation. Thus, we next tested if H2AX.2t is phosphorylated at S139 under DNA damage conditions. The chemotoxic drug bleomycin induces double strand breaks (DSBs) and increases S139 phosphorylation. We treated SUM159 cells with 10 µM bleomycin for 1 hour. There is no significant difference in the ratio of truncated to full length H2AX with or without bleomycin treatment (Fig. 3a). As expected H2AX S139ph increases upon bleomycin treatment. H2AX.2t S139ph is not observed with or without treatment despite S139 being retained on its C-terminus (Fig. 3b-c). Thus, H2AX truncation impairs γH2AX signaling. The absence of S139ph on H2AX.2t is likely explained by the loss of most of the C-terminal side of the ATQA**S**QEY motif recognized by ATM/ATR.

**Fig. 3.**
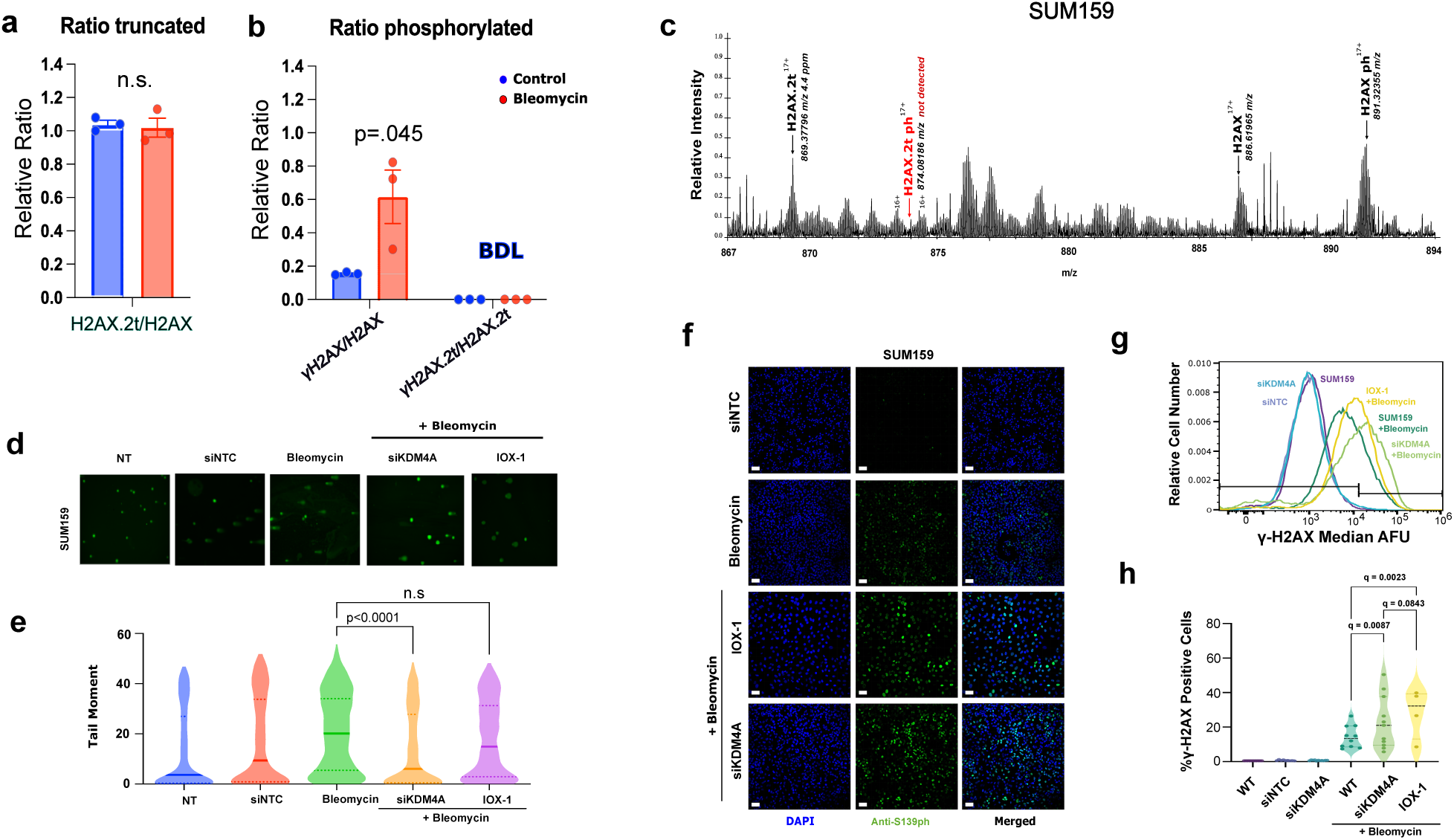
Truncation of Histone H2AX attenuates the DNA damage response. **a**, In SUM159 cells approximately 50% of H2AX exists as H2AX.2t. This ratio is unaffected by bleomycin treatment, (N=3). **b**, Bleomycin treatment induces phosphorylation of full length H2AX generating γ-H2AX. Serine 139 phosphorylation is not detected on H2AX.2t with or without bleomycin treatment, (N=3). **c**, MS1 spectrum averaged from 39.63-43.25 minutes reveals abundant γH2AX^+17^ at 891 *m/z* but no evidence of phosphorylated H2AX.2t^+17^ at 874 *m/*z. Phosphorylated H2AX.2t is not observed at other charge states, retention times, or conditions (not shown). **d**, SUM159 cells with KDM4A knockdown reduces accumulated damage by COMET tail length; however, N-Octyl IOX-1 only exhibits a minor reduction in tail length. **e**, Quantification of comet tail moments per condition tested. **f**, IF images reveal increased γH2AX signaling (S139 phosphorylation) with KDM4A knockdown in SUM159 cells. (S139ph: #9718S, AF488: #22C0617) **g**, Representative histogram displaying various conditions analyzed for γH2AX. Both N-Octyl IOX-1 and KDM4A knockdown enhance γH2AX signal in a bleomycin background. (S139ph: #05636, Phospho P53:#9286) **h**, Quantitation of γH2AX by flow cytometry shows increased signal with both N-Octyl IOX-1 treatment and KDM4A knockdown relative to bleomycin alone. (S139ph: #05636, Phospho P53: #9286, AF647: #A21236)

### Knockdown of KDM4A increases γH2AX and reduces accumulated DNA damage

Because H2AX.2t generation depletes full length H2AX and H2AX.2t is itself refractory to S139 phosphorylation, γH2AX quantification may not be the best indicator of actual DNA damage. Thus, we used an alternative approach to assess DNA damage on a cellular level. The COMET assay developed by Johannson et. al, is a single cell gel electrophoresis method that allows for the visualization of DNA breaks in cells^32^. We used this technique after inducing DSB with bleomycin and with or without knockdown of KDM4A, then quantified DNA damage per condition (Fig. 3d-e). Quantifying tail moments (tail length corresponds to DNA damage), reveals a significant decrease in DNA damage in cells when expression of KDM4A is reduced. The effect on accumulated DNA damage with N-Octyl IOX-1 treatment is not statistically significant despite its strong effect on H2AX.2t (Supplementary Fig. 4). This is likely due to its broad inhibition of other class 2 demethylases that are involved in the DNA damage response and repair process^33^. Thus, we decided to repeat these conditions and measure DNA damage signaling by S139 phosphorylation.

We used IF to quantify γH2AX (Cell Signaling Technology, Cat #20E3) in WT and KDM4A knockdown cells before and after treatment with bleomycin. Contrary to the COMET results, we found that S139 phosphorylation actually increases upon KDM4A knockdown and upon N-Octyl IOX-1 treatment (Fig. 3f). We then confirmed, using a different γH2AX antibody (Sigma Aldrich #05636) and flow cytometry, that inhibition or depletion of KDM4A leads to increased levels of γH2AX. Statistically significant increases are observed for both inhibition of KDMs with N-Octyl IOX-1, q=0.023 and knockdown of KDM4A, q=.0087 (Fig. 3g-h). These results show that reducing KDM4A activity depletes H2AX.2t, restores γH2AX signaling, and support the corollary that this increases effective DNA repair. Thus, we have identified a mechanism consistent with our hypothesis that KDM4A catalyzes cleavage of H2AX, and this modulates the DNA damage response and repair system.

## Discussion

The sequence specific post-translational truncation of two C-terminal amino acids is a previously unknown mechanism of regulation of protein function and modulation of signaling. Known carboxypeptidases remove one amino acid at a time but may do so sequence specifically and progressively, leaving evidence of each step^34^. There are many examples of the functional importance of C-terminal post-translational proteolytic processing^35^. The systematic analysis of such events is challenging; however, the loss of two C-terminal amino acids is far more common than expected based on currently known mechanisms^29^. Thus, the mechanism described here could be a common means of protein regulation and signaling.

Proteases have evolved multiple times, sometimes convergently. There are diverse protease mechanisms; however, most function by some form of hydrolysis. New protein hydrolase mechanisms have been discovered as recently as 2004^36^. The Asparagine Peptide Lyases, discovered in 2010, are significantly different in that water is not required^37^. The jumonji domains of KDM proteins, such as KDM4A, are fundamentally alpha-ketoglutarate dependent dioxygenases^25^. Thus, the protease activity we observe likely occurs through a mechanism distinct from other proteases and with a novel dependency on the carboxylic acid cycle.

Although best known for their ability to remove methyl marks, evidence has emerged revealing KDMs can act as histone hydrolases, expanding the repertoire of their function^26^. In the context of H2AX.2t, a class two KDM (KDM4A), functions as a protease through its dioxygenase activity to mediate loss of H2AX’s C-terminus. This previously unknown protease activity of KDM4A suggests a new distinct function of the enzyme, and we speculate that other class 2 enzymes may work in a similar fashion. The reason KDM4A commits to one function over the other is unclear. However, it remains of interest to elucidate the contexts and mechanisms that drive proteolysis over demethylation. Therefore, not only have we identified a mechanism of H2AX truncation, but we also discover a novel function for an extensively studied enzyme family.

An unbiased investigation of the histone H2A family reveals, here, an H2AX post-translational truncation that modulates the DNA damage response. This occurs primarily by precluding or impairing important H2AX C-terminal PTMs. We implicate KDM4A in the truncation of H2AX. It is thus a key modulator of a previously undescribed mechanism that can inhibit the canonical DDR (Fig. 4). This discovered mechanism fills a crucial gap in our understanding of these systems. This mechanism is active in multiple cellular contexts and in various tumors and tissues. Our model predicts that in low KDM4A backgrounds DDR signaling is functional, γH2AX will be relatively high, and thus accumulated DNA damage will be low (Supplementary Fig. 7a). Conversely, when KDM4A is high the DDR signaling will be insufficient and γH2AX will be underrepresented at actual damage events. This results in an accumulation of unresolved DNA damage (Supplementary Fig. 7b). This discovery implies that our current reliance on γH2AX as a marker of DNA damage is insufficient.

**Fig. 4.**
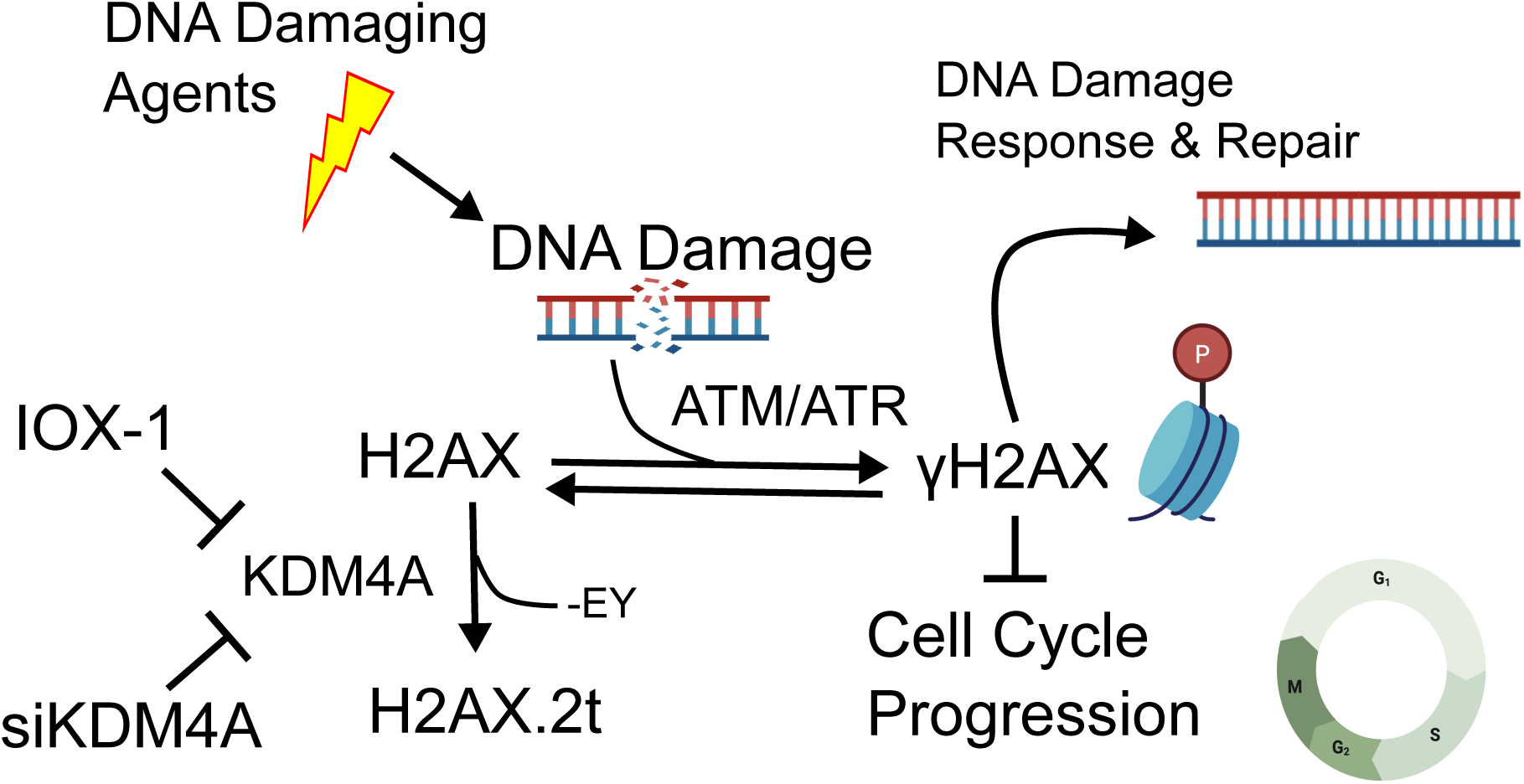
KDM4A truncation of H2AX precludes the canonical γH2AX signal and thus presents an alternate mechanism for cells to modulate the DNA damage response. A circuit model diagram of the mechanism by which KDM4A mediated truncation of H2AX supersedes the canonical DNA damage response. In this model the relationship between H2AX, H2AX.2t, and the DNA damage response is dependent on KDM4A expression. Upon DNA damage, ATM/ATR phosphorylates H2AX. γH2AX prevents cell cycle progression and initiates the DNA damage response and repair process. KDM4A cleaves H2AX, yielding H2AX.2t. Both N-Octyl IOX-1 treatment and KDM4A knockdown reduces the consumption of H2AX and the concomitant production of H2AX.2t.

The reduced substrate availability for ATM/ATR activity also inhibits the next steps of recruiting repair factors through protein-protein interactions. The DDR complex Mediator of DNA Damage checkpoint 1 (MDC1) has strong selectivity for a Tyr or Phe residue at the +3-position yielding a p-S-X-X-Y/F consensus binding motif^38,39^. Histone H2AX possesses the “p-S-X-X-Y/F” motif which MDC1 binds, on its extreme C-terminus. This motif is disrupted by the clipping of H2AX to the truncated form. Thus, not only is H2AX.2t S139 refractory to ATM/ATR phosphorylation, but the binding of DDR factors to the consensus sequence is further compromised by the loss of Y142.

The inability of H2AX.2t to respond to DNA damage suggests that it leads to a DNA damage checkpoint escape route. The onset of checkpoint proteins serves as a crucial “brake” to prevent the propagation of damaged genetic material to daughter cells. Checkpoint adaptation is a phenomenon that allows cells to overcome arrest signals induced by DNA damage checkpoints. This leads to cell cycle progression in the presence of unrepaired or partially repaired DNA lesions^40^. Without activation of checkpoint proteins, there is no delay in the cell cycle and defective cells can continue to divide. In some cases, such as the rapid proliferation and death of fibroblasts during wound healing, this may provide a fitness advantage. More generally, the capacity to reduce or completely turn off genome maintenance when not needed provides crucial energy conservation. Controlled imprecision is even exploited for a fitness advantage in certain biological processes^41^. However, when this system is aberrantly activated it will likely lead to genetic instability, mutation, and cancer. Thus, organisms must maintain a finely tuned equilibrium between energy utilization, genome stability, and genetic adaptability.

Jumonji KDM subfamilies (i.e. 2-8) are often dysregulated in cancers^42^. For example, KDM4 members are overexpressed in over 60% of all breast tumors and vary in expression based on tumor type and grade^43^. Due to this, KDM-specific therapies have emerged for the treatment of cancers; however, a more complete mechanistic understanding of KDM4A biology will inform effective use. For example, survival prognosis based on KDM4A is strong but contradictory between adjuvant and neoadjuvant therapies (Supplementary Fig. 5). KDM4A’s role in cancer is thought to be due to it altering DNA repair pathways and disrupting genomic stability^44^. The model we propose here mechanistically explains these previous observations. This elucidated mechanism should be useful in future applications in precision medicine and combinatorial therapies.

## Conclusions and Future Directions

We show here that KDM4A proteolysis activity represses H2AX S139 phosphorylation and prevents activation of the DNA damage response. By blocking the activity of KDM4A, we observe a decrease in H2AX truncation, restoration to DNA repair pathway signaling, and a decrease in actual DNA damage. This novel mechanism supersedes the known genome maintenance pathways. This discovery elucidates the mechanism of differential sensitivity of cells and tissues to DNA damage. However, it also reveals many new questions and defines new gaps in knowledge for future inquiry. For example, other histones may be cleaved in a similar manner. We have indeed observed this for H2AZ.1 but not H2AZ.2 (Supplementary Fig. 2). Additionally, while we observe this event in many patient tumors and normal tissues, we have just begun to speculate about its role in cancer development and in normal biological processes. We suspect that inhibition of KDM4A will profoundly affect tumors that express H2AX.2t, although we have not yet tested this. We have not yet demonstrated its clinical utility but expect that it will be highly beneficial in precision medicine as we have shown that this directly affects known therapeutic mechanisms. Targeting epigenetic enzymes like KDM4A as an adjuvant to the current developed treatments is promising, particularly those leveraging differences in the DNA damage response. This may slow the mutation rate and evolution of tumors, thus preventing therapy resistance and cancer recurrence. This also suggests new opportunities for prognostic stratification in cancer. Additional potential clinical applications include a better understanding of wound healing and related proliferative processes. Within the realm of bioengineering, genome editing relies on the DDR and the ability to manipulate the efficacy of this system may be essential to higher fidelity applications. Overall, this newly discovered mechanism substantially changes our understanding of the DDR and fundamental aspects of cell biology. We report here the discovery of this pathway and the validation of the essential mechanisms.

## Materials and Methods

### Alkaline Comet Assay

Comet assays were conducted as previously described^45^. Cells were resuspended to 1 x 10^5^ cells/mL and mixed with 1% low-melting agarose (Cat #4250-050-02, R&D Systems) at a 1:10 ratio and plated on 2-well comet slides (Cat #4250-200-03, R&D Systems). Cells were then lysed overnight and immersed in alkaline unwinding solution as per manufacturer’s protocol (Trevigen). Slides were stained with SYBR green (Cat #SS11494, Invitrogen) and fluorescence microscopy was performed at 10X magnification using the Keyence BZ-X800 microscope and analyses of comet tails were performed using the Comet Assay IV software (Instem).

### Immunofluorescence Assays

The cells were plated on µ-Slide 2 Well (Cat #80286, Ibidi) at a density of 9.0 x 10^5^ and for some experiments were treated with either N-Octyl IOX-1 (Cat #530537001, Fisher), or bleomycin (Cat #B8416-15UN, Sigma Aldrich). Any plasmids used for gene knockdown were used at 10uM for 24-48 hours prior to imaging. The cells were washed with Phosphate Buffered Saline (PBS), fixed with 4% Paraformaldehyde (PFA), then permeabilized and blocked with 0.5% Triton X-100 and 0.5% Bovine Serum Albumin (BSA) in PBS for 1 hr. The cells were stained with primary antibody overnight at 4C, then washed with PBS several times and stained with Alexa Fluor 488-conjugated secondary antibody (Cat #22C0617, Millipore) for 1 hr. Nuclei were counterstained with 4′,6-diamidino-2-phenylindole (DAPI). Images were acquired through Nikon A1-R confocal microscope. The exported images were analyzed by the Integrated Microscopy Core at BCM, and representative images were edited for appearance and did not affect quantitation. Slides were stained for the following targets-H2AX.2t (Cat #PP145 or #PWP201, Cell Signaling Technology), S139 phosphorylation (Cat #9718S, Cell Signaling Technology) then AF488 staining was used prior to imaging (Cat #22C0617, Millipore). Note all H2AX.2t reagents (antibodies or plasmids) listed within our methods were gifted by Cell Signaling Technologies in support of this work. These items are not yet commercially available.

### Western Blot

To lyse cells, a 10X lysis buffer was used (Cat #9803, Cell Signaling Technology) before lysates were sonicated on ice for 30 s three times with a 30 s pause between each sonication then frozen at –80C. Total protein concentration was determined using a Pierce BCA assay kit (Cat #23250, Thermo Scientific). Protein (30 μg total per well) was loaded on a 12% Mini-PROTEAN TGX gel (Cat #4561045, Biorad) and then transferred to a 0.2 μm nitrocellulose membrane using wet transfer at 25 V for 3 m. Membranes were blocked in 5% BSA (Cat #BP1600-100, Fisher) diluted in Tris Buffered Saline with Tween (TBST) (Cat #IBB-188-1L, Boston Bio Products). Membranes were probed with primary antibody overnight at 4C and secondary antibodies in 5% nonfat dry milk for 2 h at room temperature the following day then imaged by Biorad chemidoc. Primary antibodies used include H2AX.2t (Cat #PWP201 and #PWP9208, Cell Signaling Technology), S139 phosphorylation (Cat #9718S, Cell Signaling Technology), GAPDH (Cat #14C10, Cell Signaling Technology), and β-Actin, (Cat #4967S, Cell Signaling Technology). All blots were stained with Goat anti Rabbit Secondary antibody (Cat #32460, Invitrogen).

### Cell culture

HEK293T, MCF7, WI38 and IMR90 cells were maintained in Dulbecco’s Modified Eagle’s Medium (DMEM) (Cat #11965118, Thermo Fisher) supplemented with 10% Fetal Bovine Serum (FBS) (Cat #A52567-01, Thermo Fisher) and 5% penicillin-streptomycin (Cat #15070063, Thermo Fisher) and incubated at 37 °C, 5% CO2, and the required O2 concentration.

SUM159 cells were maintained in Ham’s F-12 media (Cat #11765054, Thermo Fisher) supplemented with 10% FBS and 5% penicillin-streptomycin and incubated at 37 °C, 5% CO2, and the required O2 concentration.

Peripheral blood mononuclear cells (PBMCs) and Activated T cells were provided by the laboratory of Dr. Cliona Rooney at Baylor College of Medicine. Before obtaining, cell lines were maintained in a 37-degree incubator with 5% CO_2._

### Histone Isolation

Nuclei from SUM159 cells were isolated and sulfuric acid extraction was used to produce a crude histone extract. The histones were further purified and separated by family by off-line reverse phase HPLC as described in Holt *et al*^46^. After offline separation, H2A fractions were collected and combined directly into 200 µL autosampler vials that are compatible with on-line HPLC. Combined samples were then concentrated to dryness with a vacuum centrifuge concentrator (Savant™ SPD131 SpeedVac, Thermo Scientific, Waltham, MA).

### Online HPLC

Online HPLC further separated H2A fractions on a C3 reverse phase column which were then analyzed with an Orbitrap Fusion Lumos (Buffer A: 2% ACN, 0.1%FA, Buffer B: 98% ACN, 0.1% FA). A multi-step gradient at 0.2 µl/min was used: 28% B to 30% B in 55 minutes, 30% B to 98% B from 55 minutes to 60 minutes, holding at 98 % from 60 to 65 minutes.

### MS/MS

Nanospray Flex source was used for ionization with 1700 V in positive ion mode and standard pressure mode. Data acquisition of intact protein was performed by an Orbitrap Fusion Lumos. MS1 settings of this 75 minute method used the orbitrap detector with an 120,000 resolution setting, an automatic gain control (AGC) target of 8.0e5, 200 ms maximum injection time, 3 microscans, profile data type, positive ion mode, and quadrupole isolation. For MS2 analysis the orbitrap was used with an 120,000-resolution setting, an AGC Target of 1.0e6, 250 ms Maximum Injection Time, and 3 microscans.

### MS Data Analysis

Data analysis was performed manually by assigning recorded masses to expected masses of H2A variants. Because the masses of all interested PTMs and variant sequences were known prior to MS analysis, a mass list was generated as a reference to identify the mass of H2AX and C-terminal phosphorylation’s.

### Plasmids

SiRNA Transfections were performed with Dharmacon siRNA pools for non-targeting control, KDM4A, 4B, 4C, 5A, 6B ( Cat #D-001810-10-20, Cat #M-004292-01-0005, Cat #M-004290-01-0005, Cat #M-004290-01-0005, Cat #M-004293-02-0005, Cat #M-020709-00-0005, Cat #M-003297-03-0005, Cat #M-023013-01-0005) at 10uM for 24-48 hours. H2AX.2t plasmid (Cat #60738) was gifted by Cell Signaling Technology. Lipofectamine 2000 (Cat #11668500, Thermo Fisher) was used to introduce plasmid into cells and cells were analyzed by IF or western blot for H2AX.2t expression 48 hours later.

### Human Tissue Microarray

TMAs were constructed at the Baylor College of Medicine Human Tissue Acquisition & Pathology (HTAP) core. HTAP provided TMA slides of human malignancies and performed IHC staining with H2AX.2t antibody followed by imaging. All slides were stained with H2AX.2t #PWP9119 at 1:250 prior to imaging.

### Flow Cytometry DNA Damage Marker Quantification

Bleomycin treated cells were harvested and centrifuged at 1000xG for 3 minutes in the cold and fixed using 2% PFA in PBS for 15 minutes on ice. After fixation, cells were permeabilized using Triton X-100 0.05% in PBS on ice for 15 minutes and then blocked using filtered BSA 5% in PBS for 30 minutes on ice. Primary antibody incubation was done on a bench rotator in the dark for 1 hour. For γH2AX detection a 1:750 dilution of the anti-phospho-Histone H2A.X (Ser139) antibody (Cat #05-636, Sigma-Aldrich) was used, and for phosphorylated p53 detection a 1:1000 dilution (765 ng/ml) of the Phospho-p53 (Ser15) antibody was used (Cat #9286, Cell Signaling Technology). A 1:1000 dilution (2mg/ml) of the Alexa Fluor 647 secondary antibody was used for fluorescent labeling (Cat #A-21236, Thermo Fisher) and cells were incubated on a bench rotator in the dark for an additional hour. All antibodies were diluted in eBioscience Flow Cytometry Staining Buffer (Cat #00-4222-26, Thermo Fisher). Prior to flow cytometry analysis, cells were resuspended in Flow Cytometry Staining Buffer.

### In vitro Activity Assay

In vitro demethylase assays were performed in a buffer containing 25 mM Tris-HCl pH 7.5, 20 mM NaCl, 0.005% (v/v) Triton X-100, 1 mM tris(2-carboxyethyl)phosphine (TCEP), 250 μM sodium ascorbate, 250 μM α-ketoglutarate, and 25 μM ferrous ammonium sulfate. Recombinant human KDM4A (produced by the Baylor College of Medicine Recombinant Protein Production Core Facility) was used at a final concentration of 1μM. H2AX.2t (Cat #PWP9119, Cell Signaling Technology) was used to detect H2AX.2t levels in various conditions of the reaction. Substrates included recombinant histone H3K36me3 (Cat #31580, Active Motif) at 2 μM, or H2AX peptide (sequence H-CKATQASQEY-OH, Innopep) at 200 μM. KDM4A and substrate were combined in demethylase buffer and incubated at room temperature for 2 h. Where indicated, ferrous ammonium sulfate was omitted to serve as negative controls. Demethylation reactions were directly analyzed by dot blotting at the indicated substrate concentrations.

### Alpha Fold Prediction Software

The protein-protein interaction maps and structure were predicted using Alphafold2 in ChimeraX 1.3. Uniprot sequences O75164 and P16104 were used.

## Data availability

Raw proteomics data is available in the following repository: (Reviewer Access)

## Statistical Analysis

All methods unless specified otherwise, utilize Welch’s two tailed t-test. All error bars represent standard error of the mean.

Flow cytometry data was analyzed in the following manner: gross cell and singlets populations were gated out prior to defining the gH2AX High/Low gate.

Two sets of analysis were performed to quantify High/Low DNA Damage:

0.50WT Gating: the gH2AX high gate was set Arbitrarily to indicate 0.50% of SUM159 untreated cells to be gH2AX high.

This is the method chosen in the final figure due to the higher reproducibility and lower chance of human error.

Peak Gating: The gate was manually set at the approximate apex of the SUM159+Bleo sample curve. This decision was made after comparing the SUM159 untreated and SUM159+Bleo treated samples and observing this point to be where the difference between the curves was highest.

While this method shows more significance, it’s less reproducible as it is based on user observation. It would be interesting to include in supplemental information if desired.

The chosen histogram plot for the final figure is indicated with several red ‘XXXXX’ in the figures containing the histogram pots for all replicates. Please include the full detailed plots for all replicates in the supplemental information. This might help us avoid reviewers questioning the reproducibility of flow cytometry.

### Flow cytometry statistical analysis

For the 0.50WT gating data, all samples passed a Normality test. For the Peak Gating data, all but the SUM159 dataset passed a Normality Test.

A One Way ANOVA test was chosen for analysis and ONLY the datasets depicted in the graph were included in the analysis. Excluded datasets, deemed biologically irrelevant, were not included.

Final analysis was done with an FDR correction, with the assumption of a p = 0.05 (5% chance of false positives). Resulting in adjusted p-values, q-values, that indicate significance.

An alternative analysis using Tukey cannot detect significance as it is more stringent than FDR, since it controls for the chance of committing one or more false positives.

## Acknowledgments

This work was supported by the National Institutes of Health grants to NLY: R01GM139295, P01 AG066606, R01 CA193235, R01 AG074540, R01 CA276663, and R01 NS136375 & to SMR: R01 CA250905 and DP1 AG072751. We thank Johnathan R. Whetstine for consulting on KDM4A activity and first suggesting the modeling experiment in Fig. 2i and Supplementary Fig. 3. We thank Margaret A. Goodell for use of resources and equipment. We also thank the members of our laboratories and many colleagues throughout the scientific community for additional critical feedback and advice. The H2AX.2t antibodies used are a kind gift from Cell Signaling Technology.

## Author Contributions

F.M.J., M.V.H & N.L.Y. conceived of the experiments. F.M.J., M.V.H, R.D., S.M.R., D.R.R. & N.L.Y. planned the experiments. F.M.J., M.V.H, J.M.J, L.Z., A.G.B., P.J.C., & S.I.A. carried out the experiments. F.M.J., M.V.H, J.M.J, L.Z., A.G.B., P.J.C., S.I.A., R.D., S.M.R., D.R.R., N.L.Y. contributed to the interpretation of the results. F.M.J. & N.L.Y. wrote the manuscript. All authors provided critical feedback and helped shape the research, data analysis, interpretation, and manuscript.

