## Supplementary Data for "H2AX C-Terminal Dipeptide Truncation: A Master Switch of the DNA Damage Response"

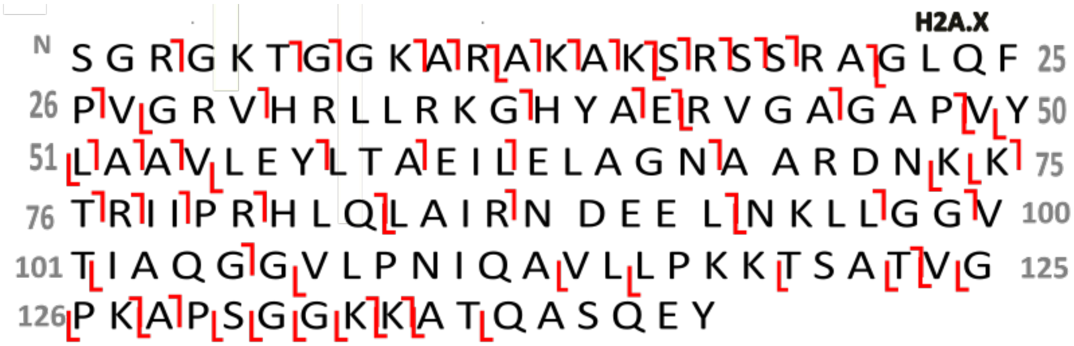


**Supplementary Fig. 1 | Ion map showing the fragmentation pattern of full length H2AX in SUM159 cells.**

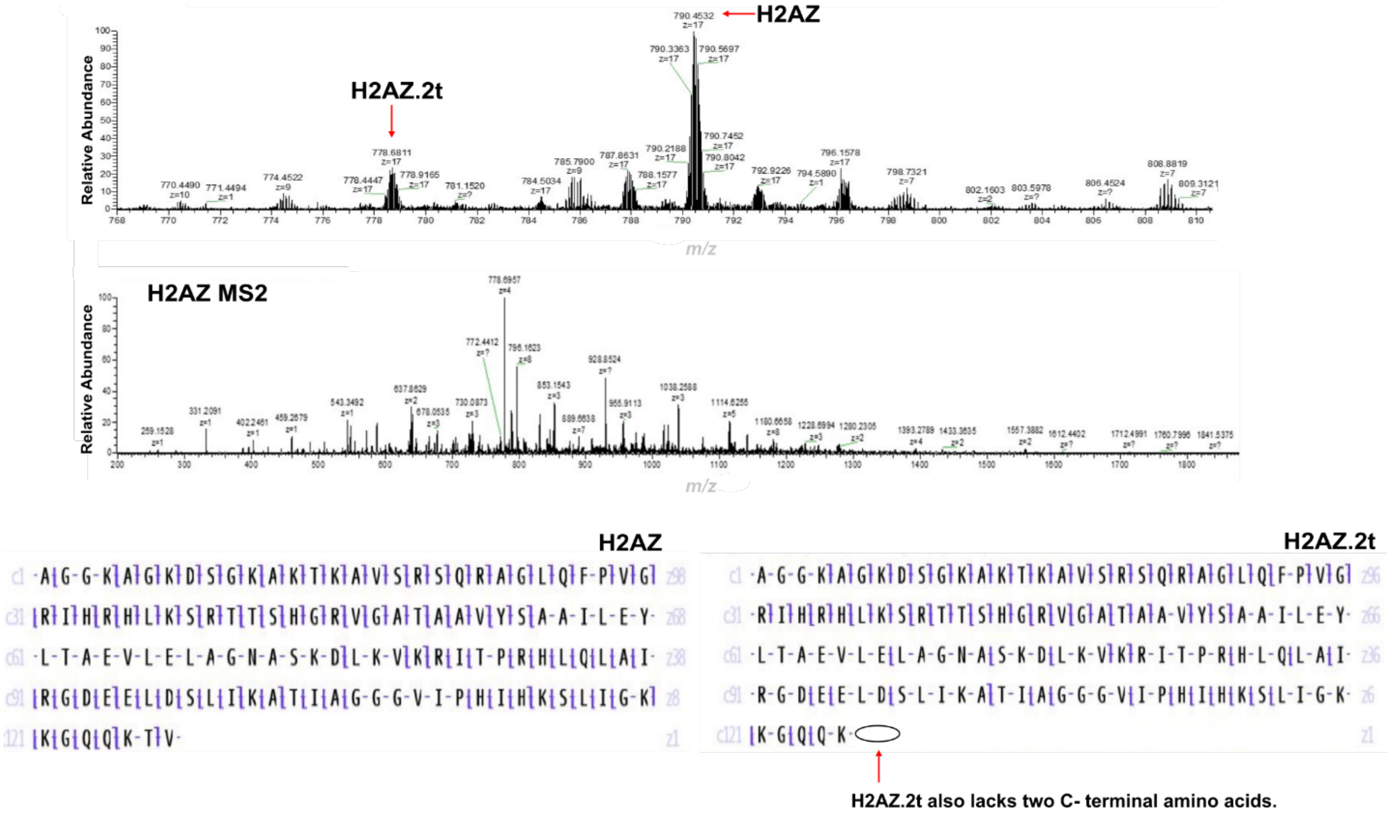

**Supplementary Fig. 2 | Histone variant H2AZ.1 is also cleaved by two amino acids on its C-terminus.**

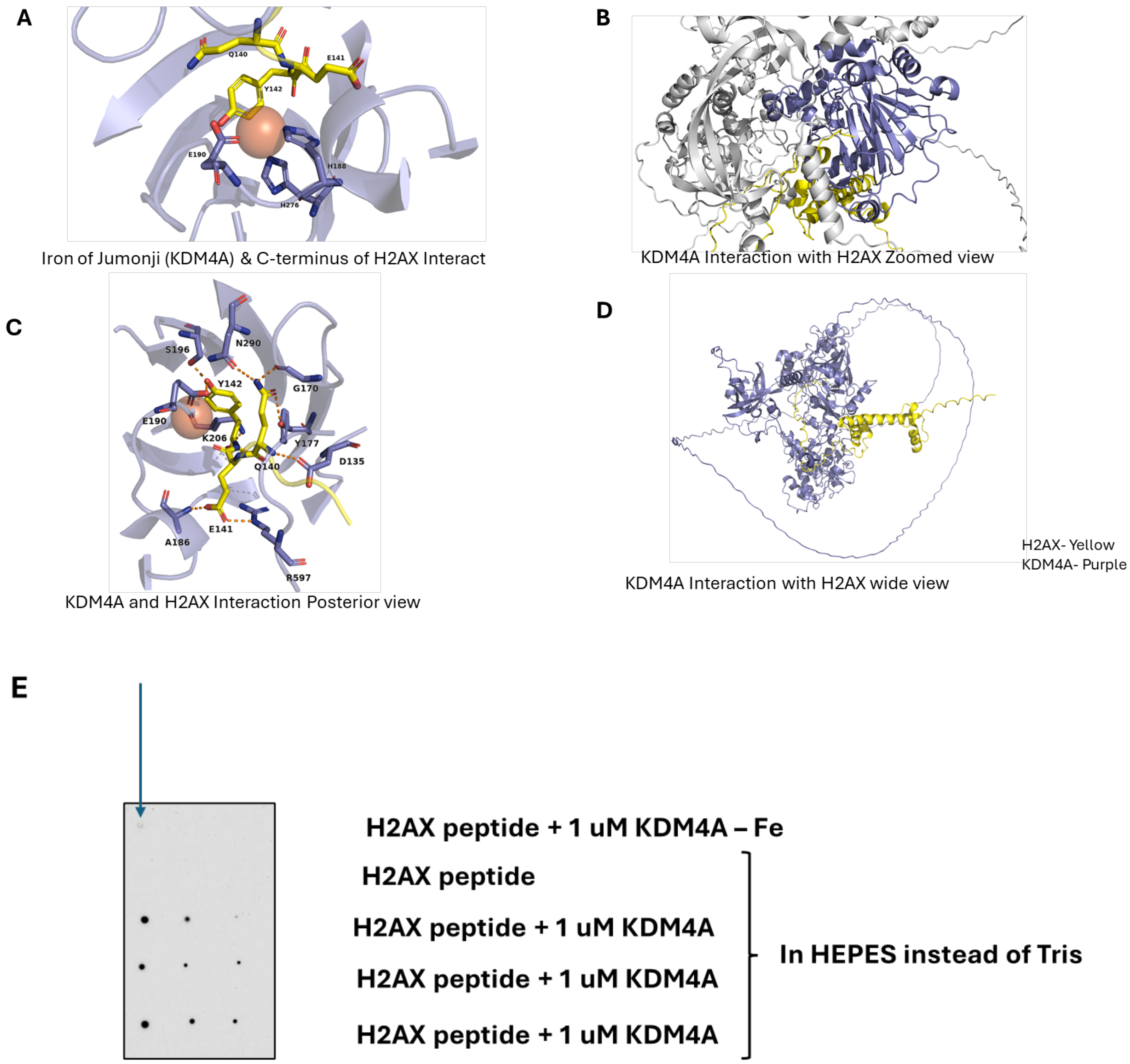

**Supplementary Fig. 3 | Predicted Alpha Fold structure of full length H2AX, KDM4A interactions, and in vitro iron sensitivity. a,** The C-terminus of H2AX is predicted to interact with the active site of the KDM4A Jumonji domains. The KDM4A contains an active site iron that is chelated by H188, H276, and E190. H2AX Y142 is predicted to directly interact with the active site iron, suggesting a mechanism of sequence specificity. This would bring the peptide bond between Q141 and E142, the site of proteolysis, in close unhindered proximity of the active site iron. **b-d)** The structured domains of full length H2AX and KDM4A are also predicted have complementary interactions in this configuration. **e**, Exclusion of iron during an in vitro reaction displays reduced activity between KDM4A and H2AX.2t (blue arrow). KDM4A and H2AX.2t activity is retained when the reaction is performed in HEPES instead of Tris buffer.

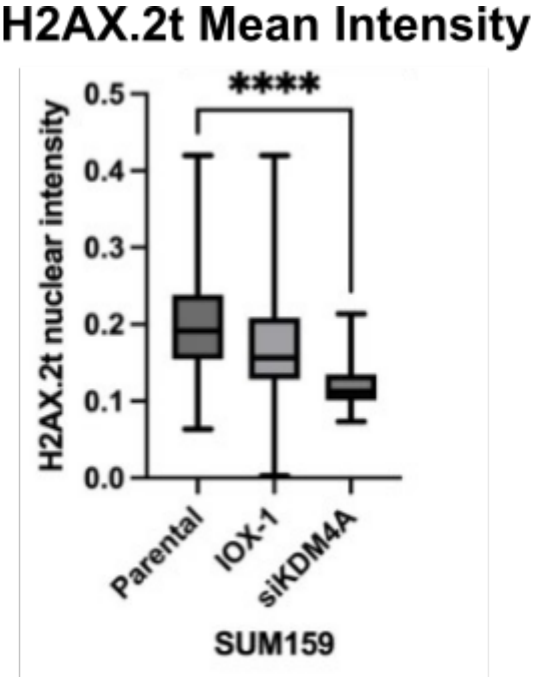

**Supplementary Fig. 4 | Mean Intensity of H2AX.2t expression in SUM159 cells.** Nuclear intensity of H2AX.2t is significantly reduced when KDM4A is knocked down in SUM159 cells.

Cohort

Cohorts: not selected

Results

**P value:** 2.9e-6

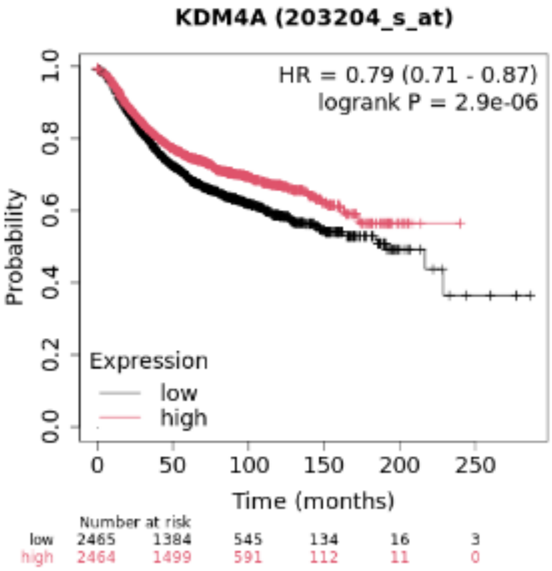


Download plot as a PDF

Upper quartile survival

| Low expression cohort (months) | High expression cohort (months) |
|---|---|
| 43 | 60 |

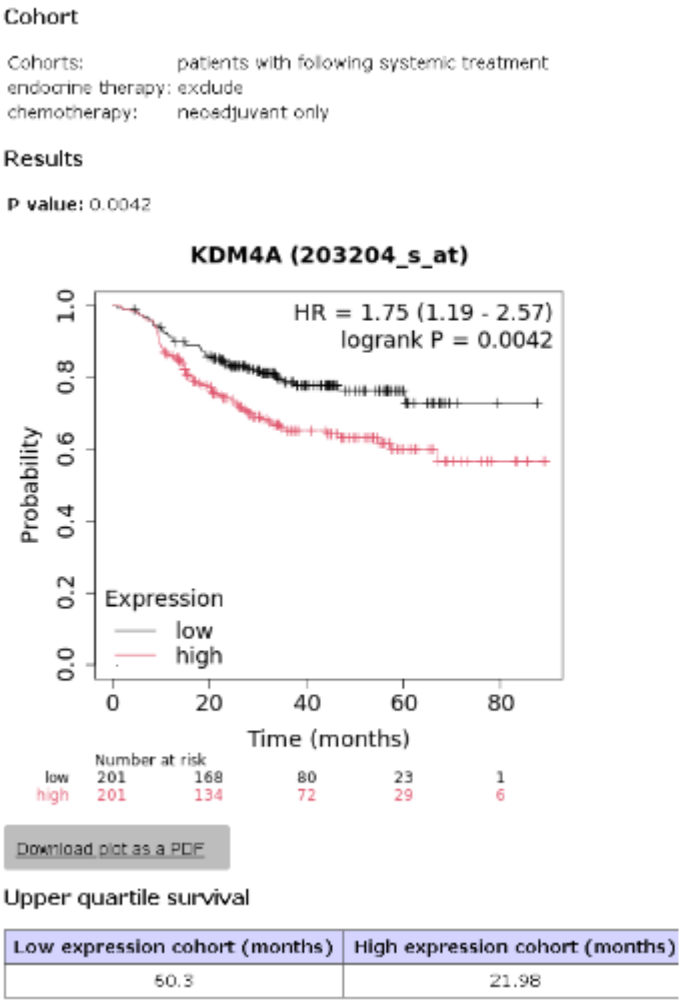


**Supplementary Fig. 5 | KDM4A Cancer Prognosis.** Above, km plot for breast cancer (all subtypes all treatments) for KDM4A is associated with better prognosis in breast cancer. Below, patients who only have neo-adjuvant chemotherapy have a worse prognosis when KDM4A is expressed.

**Supplementary Table 1 | H2AX.2t expression is confirmed in the following human tissues**

| ID | TMA | ORGAN/TISSUE | DIAGNOSIS |
|---|---|---|---|
| 60 | 5 | LYMPH NODE | PAPILLARY CARCINOMA |
| 756 | 5 | OVARIAN | SEX CORD TUMOR WITH ANNULAR TUBULES |

| | | | |
|---|---|---|---|
| 754 | 5 | OVARIAN | GRANULOSA CELL TUMOR |
| 753 | 5 | TESTICLE | GRANULOSA CELL TUMOR |
| 752 | 5 | OVARIAN | GRANULOSA CELL TUMOR |
| 738 | 5 | RETROPERITONEAL | LIPOSARCOMA |
| 720 | 5 | CERVIX | NORMAL TISSUE |
| 726 | 5 | STOMACH | GASTROINTESTINAL STROMAL TUMOR (GIST) |
| 716 | 5 | LYMPH NODE | ADENOCARCINOMA |
| 718 | 5 | OVARY | DUCTAL CARCINOMA |
| 526 | 3 | LIVER | METASTATIC ADENOCARCINOMA |
| 348 | 1 | LYMPH NODE, INTRAPAROTID | MET MALIG MELANOMA |
| 220 | 3 | MENINGES | MENINGIOMA |
| 306 | 2 | LIVER | HEPATO CELLULAR CARCINOMA (HCC) |
| 045 | 3 | DURA MATER | MENINGIOMA |
| 090 | 3 | LYMPH NODE | POORLY DIFF ADENOCARCINOMA |

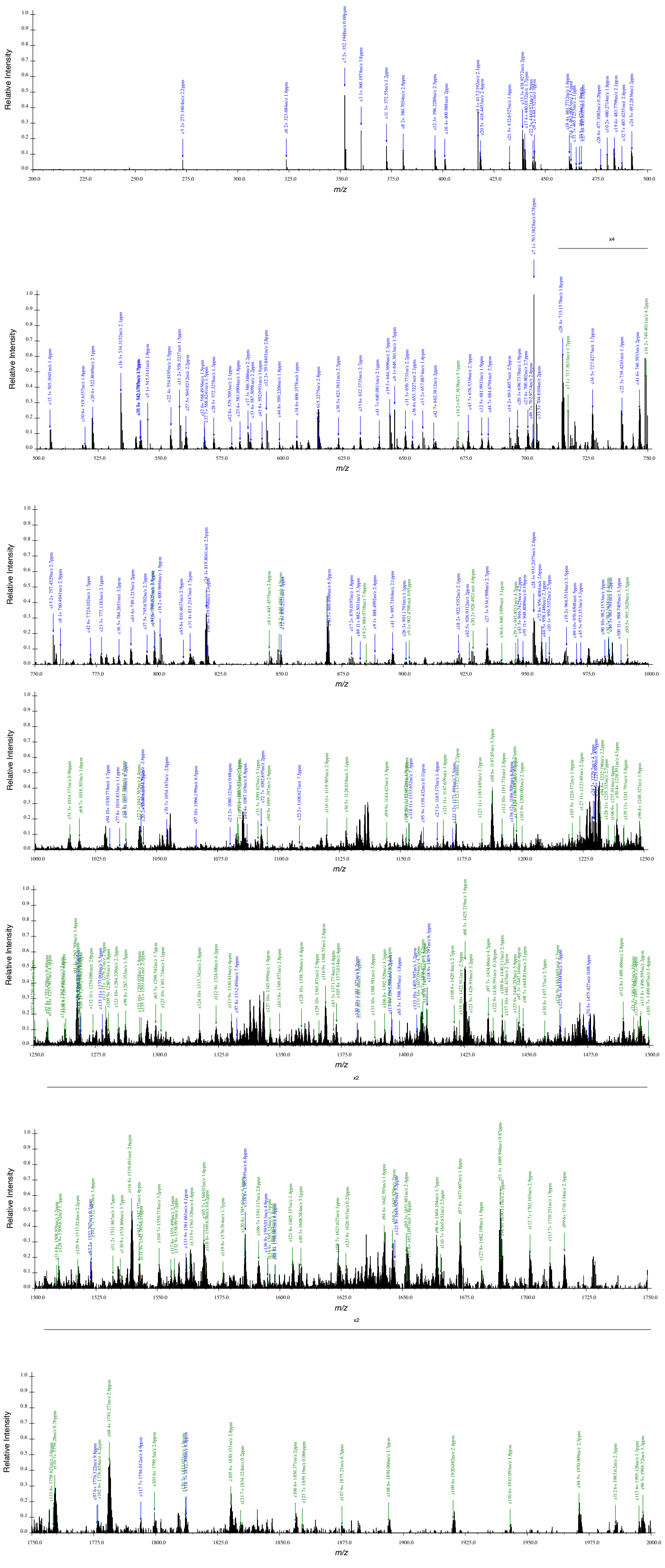


**Supplementary Fig. 6. | Supplemental Figure 6. MS2 spectra of H2AX.2t in SUM159 cells**

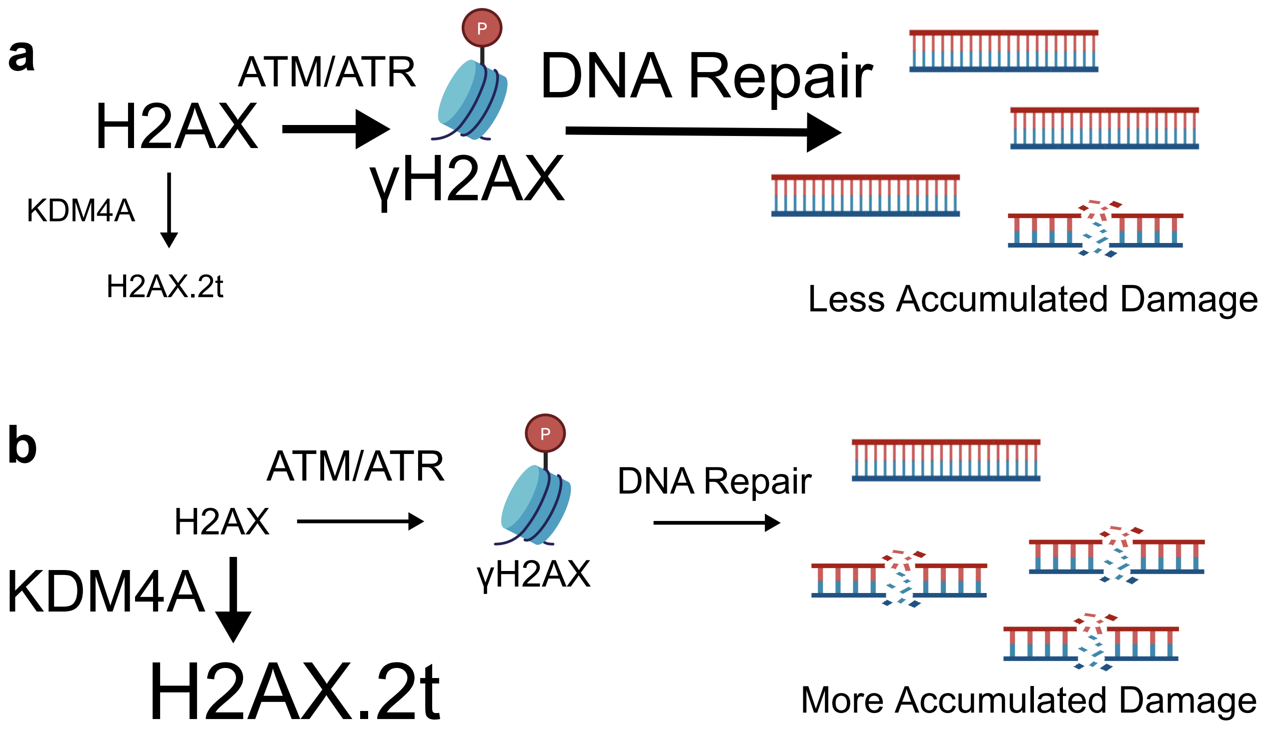


**Supplementary Fig. 7. | How KDM4A activity affects H2AX function. a,** Under normal conditions where KDM4A activity is low, there is sufficient H2AX available as substrate for ATM/ATR to signal for the DNA damage response. In this context, there is sufficient DNA repair to maintain genome stability and prevent DNA damage accumulation. **b**, When KDM4A activity is high, the pool of H2AX is depleted by conversion to H2AX.2t. Thus, there is less H2AX available as a substrate for ATM/ATR phosphorylation. This results in less DNA repair and more accumulated damage.
